# Survival of adult barn owls is linked to corticosterone levels

**DOI:** 10.1101/517201

**Authors:** Paul Béziers, Fränzi Korner-Nievergelt, Lukas Jenni, Alexandre Roulin, Bettina Almasi

**Author notes:** Shared senior authorship (Alexandre Roulin, Bettina Almasi).

## Abstract

Glucocorticoid hormones, such as corticosterone, are fundamental in the translation of external stimuli into physiological adjustments that promote the survival of an organism in face of changes in its environment. At baseline levels, corticosterone is crucial in regulating daily life metabolism and energy expenditure, whereas the acute corticosterone response promotes short-term physiological and behavioral responses to unpredictable environmental challenges. Given their different physiological effects and their role in mediating fitness components, it is still unclear whether and how inter-individual variation in baseline corticosterone levels and acute stress-response levels can affect the survival of organisms. We used 13 years of capture-recapture and dead recovery data combined with 11 years of corticosterone measurements taken on breeding barn owls (*Tyto alba*) to investigate how survival probability varies in relation to baseline and stress-induced corticosterone levels. Our study shows that males with a higher level of both baseline and stress-induced corticosterone levels have a higher probability of surviving than individuals with lower corticosterone levels. In females, survival probability was higher in individuals presenting elevated stress-induced corticosterone levels but was not significantly associated to baseline corticosterone levels. Our results suggest that in the barn owl the stress-induced corticosterone response is a better proxy of adult survival than baseline corticosterone levels. Further studies investigating the link between corticosterone levels and different fitness components as well as the environmental factors (*e.g*. weather, development conditions, disease and predation risk) leading to such endocrine phenotypes are needed to identify the costs and benefits of presenting high and low corticosterone profiles.

## Introduction

Glucocorticoids hormones, such as corticosterone, are crucial for the functioning of organisms as they contribute to regulate a number of important physiological processes, including food intake (Dallman et al., 1993; King, 1988), energy allocation (Landys et al., 2006; Sapolsky et al., 2000) and locomotor activity (Landys et al., 2006; Overli et al., 2002a). Baseline glucocorticoid levels are responsible for maintaining energy homeostasis in relation to current and predictable changes in energetic demands (Sapolsky et al., 2000). To sustain these activities, glucocorticoids follow a diel and seasonal rhythm in response to the different energetic demands associated to daily and life-history stages (Carsia and Harvey, 2000; Landys et al., 2006; Romero, 2002; Roulin et al., 2010). Glucocorticoids are also part of the adrenocortical stress response that allows the reallocation of resources to physiological functions that are essential to survival when the environment becomes unpredictably challenging (e.g. predator attack, food restriction, inclement weather). Glucocorticoids therefore play an important role in mediating trade-offs between different life-history traits and, as a consequence, could be associated to fitness-related traits (Crespi et al., 2013; Hau et al., 2010; Ricklefs and Wikelski, 2002; Zera et al., 2007). They may also play an important role in selective processes underlying the expression of these traits given that variation in glucocorticoid levels can have a heritable component (Brown and Nestor, 1973; Evans et al., 2006; Jenkins et al., 2014; Odeh et al., 2003; Pottinger and Carrick, 1999; Satterlee and Johnson, 1988). Although, baseline and acute stress-induced glucocorticoid levels can be associated with fitness components, such as reproductive success (Bonier et al., 2009a; Riechert et al., 2014; Schmid et al., 2013) and survival in adults and fledglings (Blas et al., 2007; Cabezas et al., 2007; Rivers et al., 2012; Romero and Wikelski, 2001), it is still unclear how variation in corticosterone levels affects the survival of an individual (reviewed in Bonier et al., 2009a; Breuner et al., 2008). Moreover, our knowledge about the effect of glucocorticoids on fitness traits mostly comes from studies where glucocorticoids levels have been experimentally elevated via implants and injection of glucocorticoids or with the increase of natural stressors, such as food shortage or predators (reviewed in Bonier et al., 2009a; Breuner et al., 2008; Crossin et al., 2016; Sopinka et al., 2015). In these studies, the experimental manipulation may not mimic perfectly natural situations. For these reasons, it is important to investigate survival in relation to natural variation in glucocorticoid levels (e.g. Blas et al., 2007; Cabezas et al., 2007; Romero and Wikelski, 2001; Wey et al., 2015; Wilkening and Ray, 2016).

It is commonly assumed that low glucocorticoid levels are beneficial because high and chronically elevated levels can impair the health of an individual (Bremner, 2007; Breuner et al., 2008; de Kloet et al., 1999; Martin, 2009; Sapolsky et al., 2000). In contrast, a rapid rise in glucocorticoid levels, in response to a stressful event, is thought to be advantageous, as it allows an organism to redirect energy stores from body maintenance and reproductive activities towards physiological and behavioral processes that are essential for survival (Almasi et al., 2008; Ouyang et al., 2012). However, the empirical evidence supporting these predictions is mixed with some studies having found positive, negative or no relation between glucocorticoids and fitness components (reviewed in Bonier et al., 2009a; Breuner et al., 2008). For instance, lizards (*Uta stansburiana* and *Lacerta vivipara*) with higher baseline corticosterone levels have a higher survival prospect (Comendant et al., 2003; Cote et al., 2006), whereas the opposite pattern has been reported in adult cliff swallows (*Petrochelidon pyrrhonota*) (Brown et al., 2005). In the mountain white-crowned sparrow (*Zonotrichia leucophrys oriantha*), individuals mounting higher stress-induced corticosterone responses showed higher survival (Cabezas et al., 2007; see also Patterson et al., 2014), whereas in nestling white storks (*Ciconia ciconia*) the opposite pattern was observed (Blas et al., 2007). These inconsistencies between studies could result from variation in corticosterone levels across life-history stages and their different energetic demands, which might select for different levels depending on the stage and context in which individuals are sampled (Bonier et al., 2009a). For instance, selection may favor low corticosterone levels during the early stages of reproduction but higher levels at later stages to reallocate resources from body maintenance to parental care (Bonier et al., 2009b; Magee et al., 2006; Romero, 2002). Inconsistencies may also arise because of sex-specific roles in reproduction (Bonier et al., 2007) and individual differences in life-history strategies (Angelier et al., 2011; Lancaster et al., 2008; Schultner et al., 2013). Studying the regulation of glucocorticoids in natural populations is inherently difficult as multiple environmental factors cannot be standardized, such as climatic conditions (Bize et al., 2010; Ouyang et al., 2015; Thierry et al., 2013), predation risk (Scheuerlein et al., 2001; Sheriff et al., 2009), food abundance (Jenni-Eiermann et al., 2008; Kitaysky et al., 2007), population density (Dantzer et al., 2013; Glennemeier and Denver, 2002; Meylan and Clobert, 2004) and disease outbreaks (Dunn et al., 1989; Gustafsson et al., 1994; Love et al., 2016). Moreover, corticosterone levels are a highly plastic trait that can change in relation to an all-suite of environmental factors that are also known to affect fitness and which can ultimately bias or obscure selection patterns (Bonier and Martin, 2016).

In the present study, we investigated whether baseline and stress-induced corticosterone levels are related to adult survival in a free-living population of barn owls (*Tyto alba*). We used capture-recapture and dead recovery data collected during 13 years, with corticosterone measurements collected during 11 years in breeding adults. The barn owl breeds in human-made sites, which may have profound consequences on its survival given that human presence and landscape structure around breeding sites affect corticosterone levels in nestlings (Almasi et al., 2015).

## Materiel and Methods

### Study site and species

The barn owl is a medium-sized bird of prey that lives in open rural landscapes where it hunts on small mammals. In our study area, barn owls commonly breed in artificial nest boxes fixed to the wall of barns. From the middle of February to beginning of August, females lay 1 to 2 clutches per year each comprising between 2 and 11 eggs (Béziers and Roulin, 2016). Breeding pairs in our population are rather faithful to their breeding site, 78% of pairs who stay together from one year to another stay in the same breeding site (Dreiss and Roulin, 2014). The typical lifespan of barn owls is of 4 years but individuals up to 15 years have been recorded in our population (Altwegg et al., 2007). Reproduction and survival in this population are being recorded since 1990.

The present study was conducted between 2004 and 2016 in western Switzerland (46°49’N, 06°56’E). All nest-boxes were controlled once a month from March till September to identify breeders. Additional controls were made to ring nestlings and adults, determine clutch size and number of fledglings. Adult females (n= 289 individuals) and males (n= 166) were captured at the end of the incubation stage or during nestling provisioning (Table 1). If an adult had not been ringed as a juvenile, its age was estimated from the moult pattern of wing feathers (Taylor, 1993). We distinguished females from males by their incubation behavior and by the presence of a brood patch.

**Table 1.** Number of baseline and stress-induced samples taken from 2004 to 2016 in breeding barn owls.

|  | 2004 | 2005 | 2006 | 2009 | 2010 | 2011 | 2012 | 2013 | 2014 | 2015 | 2016 |
| --- | --- | --- | --- | --- | --- | --- | --- | --- | --- | --- | --- |
| Type of corticosterone samples |  |  |  |  |  |  |  |  |  |  |  |
| *Baseline* | 56 | 77 | 47 | 16 | 30 | 30 | 120 | 17 | 36 | 82 | 130 |
| *Stress-induced response* |  |  | 40 | 23 | 29 | 37 | 169 | 21 | 57 | 105 | 159 |
| Number individuals sampled for baseline levels |  |  |  |  |  |  |  |  |  |  |  |
| *Males* | 22 | 45 | 10 | 3 | 5 | 4 | 26 | 3 | 2 | 14 | 16 |
| *Females* | 34 | 32 | 37 | 13 | 25 | 26 | 94 | 14 | 34 | 68 | 114 |
| Number individuals sampled for stress -induced levels |  |  |  |  |  |  |  |  |  |  |  |
| *Males* |  |  | 10 | 9 | 5 | 8 | 40 | 8 | 15 | 24 | 27 |
| *Females* |  |  | 30 | 14 | 24 | 29 | 129 | 13 | 42 | 81 | 132 |

### Assessment of baseline and stress-induced corticosterone levels

To assess corticosterone levels, adult barn owls were captured and submitted to a standardized capture restrain protocol. A first blood sample was taken within 3 minutes (mean: 2’36’’ ± 32’’ (SD)) after first disturbance (*e.g*. entering the barn or triggering the trap) to assess baseline corticosterone levels (n= 641 samples, Table 1). Although the increase in corticosterone level during the first 3 minutes after an acute stress has been shown to be marginal in other species (Romero and Reed, 2005; Roulin et al., 2010), we considered sampling time in our statistical analyses (Table 2). The blood sample was taken by puncturing the brachial vein and was then collected with heparinised capillaries. Blood samples were directly centrifuged, the plasma was separated and flash frozen in liquid nitrogen. Once back from the field (within less than 24 hours), the samples were stored at −20°C until analysis within the next 6 months. After having collected this first blood sample, the birds were weighed, the length of their wing measured to the nearest mm and then placed in an opaque cloth bag until a second blood samples was taken 25 minutes (mean: 23’58’’ ± 1’51’’ (SD)) after capture to measure the stress-induced corticosterone response (n= 640, Table 1). This time lapse corresponds to the peak of corticosterone response in the barn owl (Almasi et al., 2015).

**Table 2.** Parameter estimates from a linear mixed effect model for baseline corticosterone levels and stress-induced corticosterone response induced levels measured between 2004 and 2016 in breeding barn owls. The table shows the fixed effects and estimates used to predict corticosterone values in our multistate survival models. The estimates (95% Bayesian credible intervals) of predictors are based on the draw of 50’000 random values from the joint posterior distribution of the model parameters. The year and identity were added in the models as random terms to correct for pseudo-replication. Predictors with a significant effect on corticosterone levels are written in bold. The baseline estimates are based on 641 samples taken from 361 individuals measured in 11 different years, whereas the stress-induced corticosterone is based on 640 samples taken from 341 individuals measured in 9 different years (Table 1).

| Fixed effects | Baseline model<br>estimates<sup>†</sup> (95% CrI) | Stress-induced model<br>estimates<sup>†</sup> (95% CrI) |
| --- | --- | --- |
| Intercept | -0.28 (-0.53 – -0.03) | -0.22 (-0.48 – 0.03) |
| Date of sampling | **-0.34 (-0.42 – -0.26)** | **-0.23 (-0.3 – -0.16)** |
| Hour of sampling | 0.07 (-0.01 – 0.15) | **0.07 (0.004 – 0.14)** |
| Blood sampling time* | **0.1 (0.02 – 0.18)** | **0.11 (0.03 – 0.18)** |
| Sex** | 0.06 (-0.07 – 0.19) | 0.06 (-0.07 – 0.18) |
| Body mass | **0.1 (0.02 – 0.18)** | **-0.18 (-0.29 – -0.08)** |
| Brood size | 0.06 (-0.01 – 0.12) | 0.01 (-0.06 – 0.08) |
| Breeding stage*** | **0.23 (0.14 – 0.32)** | **0.25 (0.16 – 0.33)** |
| Random effects | Variance ± sd | Variance ± sd |
| Year | 0.03 ± 0.16 | 0.04 ± 0.19 |
| Individual identity | 0.04 ± 0.20 | 0.05 ± 0.23 |
| Residual variance | 0.16 ± 0.41 | 0.14 ± 0.38 |

We indicate the variance explained by the random variables (individual identities and year) and residual variance (± sd).
<sup>†</sup> Estimates based on standardized data
\* latency between time(sec) of capture and blood sampling for baseline and stress-induced model
\*\* Difference from female to male
\*\*\* Difference from offspring provisioning to incubation

### Total corticosterone assay

Plasma corticosterone was extracted with dichloromethane and determined with an enzyme immunoassay (Munro and Stabenfeldt, 1984; Munro and Lasley, 1988) following (Müller et al., 2006). Ten microliters of plasma were added to 190μl of water and extracted with 4ml of dichloromethane. The solution was mixed for 30 minutes on a vortex machine and then incubated for 2 hours. After separating the water phase, the dichloromethane was evaporated at 48°C and corticosterone was resuspended in a phosphate buffer. The dilution of the corticosterone antibody (Chemicon; cross-reactivity: 11-dehydrocorticosterone 0.35%, progesterone 0.004%, 18-OH-DOC 0.01% cortisol 0.12%, 28-OH-B 0.02% and aldosterone 0.06%) was 1:8000. We used Horseradish Peroxidase (HRP) (1:400 000) linked to corticosterone as enzyme label and 2,2’-azino-bis(3-ethylbenzothiazoline-6-sulphonic acid) (ABTS) as substrate. We determined the concentration of corticosterone in triplicate by using a standard curve run in duplicate on each plate. If the corticosterone concentration was below the detection threshold of 1ng/ml the analysis was repeated with 15μl or 20μl plasma. If the concentration was still below the detection limit (1 baseline sample), we assigned to the sample the value of the assay detection limit (1ng/ml). Plasma pools from chicken with a low and high corticosterone concentration were included as internal controls on each plate. Intra-assay variation ranged from 3 % to 14 % and inter-assay variation from 7 % to 22 %, depending on the concentration of the internal control and the year of analysis.

### Survival Analyses

We used a multistate capture-recapture and dead recovery model to estimate true survival of barn owl adults in relation to plasma baseline and stress-induced corticosterone levels (Lebreton et al., 2009). The model allowed us to estimate the probability of recapturing an individual given it is alive and did not permanently emigrate from the study area at time *t* (*p*), and its survival from time *t* to time *t* + 1, (ϕ). We modelled survival probability (ϕ) in relation to different covariates including age (quadratic effect) and sex in interaction with corticosterone levels using the logit-link function.

To predict survival in relation to corticosterone we had to model the missing corticosterone measurements because corticosterone cannot be measured when individuals are not captured. Our first approach was to integrate a corticosterone model within the survival model (this would have corresponded to the multiple-imputation method from Little & Rubin (2002)). However, the model failed to converge and we instead fitted two separated normal linear mixed models (one for baseline, and one for stress-induced corticosterone) in R using the function *lmer* from the package lme4 (Bates et al., 2015). For the baseline corticosterone model, we included sex, body mass, sampling date (*i.e*. Julian date) and timing (*i.e*. hour) as well as blood sampling time (*i.e*. time lapse between capture and blood sampling) as predictors (Table 2). We also added brood size as a measure of reproductive investment and stage at which individuals were sampled, *i.e*., during the incubation or offspring provisioning period. All variables were normalized ((x-mean(x))/(2 × standard deviation(x))) and the identity of the individuals and the year of sampling were added as random factors. Baseline corticosterone was log-transformed to obtain a normal distribution, whereas stress-induced corticosterone was already normally distributed without any transformation. The same variables were used to model stress-induced corticosterone levels (Table 2). From these two models we used the expected corticosterone values for every individual and year given a mean value for all other covariates. By doing so, we obtained expected corticosterone values for the years when individuals were not captured and we corrected the measured corticosterone values for the hour of day, sampling time, date, mass, brood size and breeding stage (incubation or offspring provisioning). This has the advantage to obtain corticosterone values that are comparable between individuals and years. In average 62% of individuals for baseline and 72% for stress-induced corticosterone were sampled in all years between first and last capture. The number of baseline and stress-induced samples per individual ranged from 1 to 11 (mean: 1.8 ± 1.32) and from 1 to 11 (mean: 1.92 ± 1.54), respectively.

Recapture probability was modeled in relation to the number of available nest boxes in our study area (*i.e*. the number of nest boxes varied from 124 to 274 between 2004 and 2016). We modeled the probability of recovering a dead individual as constant throughout the study period. The capture-recapture/recovery matrix started with individual at age 0 (*i.e*. year of birth) and included only individuals from which we had at least one corticosterone measurement as adult. As we only included juveniles who were recaptured as adults in our analyses we might have overestimated juvenile survival (survival from age 0 to 1). Due to logistic constraints sample sizes varied between baseline and stress-induced corticosterone levels and number of individuals used for baseline and stress-induced models. For baseline, 63 males and 119 females ringed as juveniles and 129 females and 50 males ringed as adults were sampled. From these 361 individuals, 25 were recovered dead during the study period (mean ± SD age recovery: 3.7 ± 2.36) and 251 individuals gave 936 recaptures (Figure 1, mean ± SD age recapture: 2.94 ± 2.04). For stress-induced corticosterone we sampled a total of 68 males and 107 females ringed as juvenile and 46 males and 120 females ringed as adults. From these 341 individuals, 13 were recovered dead during the study period (mean ± SD age recovery: 3.38 ± 1.93) and 222 individuals gave 807 recaptures (Figure 1, mean ± SD age recapture: 2.77 ± 2.07). The age of the oldest individual recovered dead was 9 years and of the oldest individual recaptured alive was 12 years (Figure 1).

**Figure 1.**
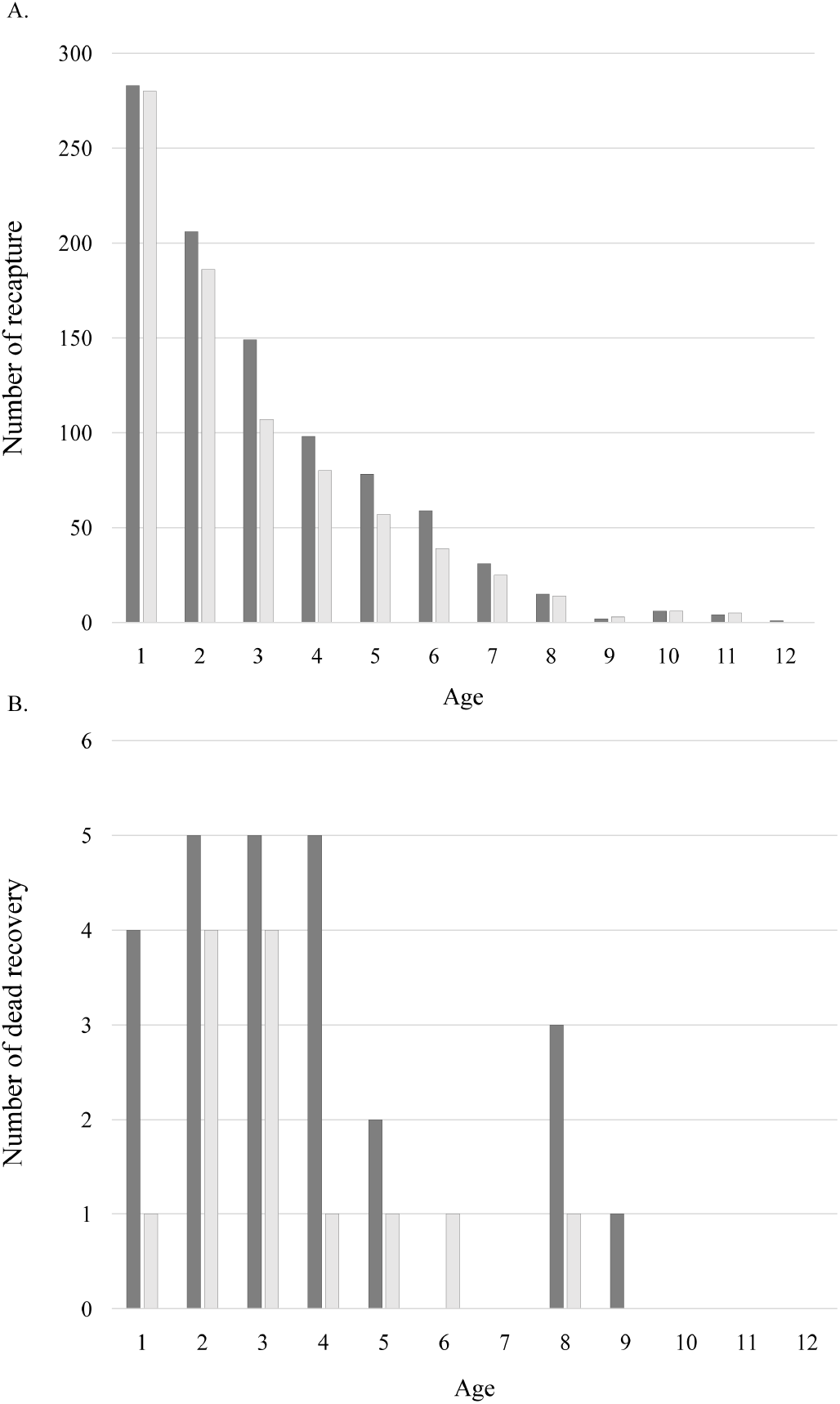
Number of (A) live and (B) dead recapture in relation to age. The dark grey and light grey bars represent the data used to estimate survival of adult barn owls in relation to baseline and stress-induced CORT levels.

All live recapture and dead recovery data from birds ringed in or out of our area were extracted from the European Union for Bird Ringing (EURING) data bank and were thus not restricted to our study area.

The capture-recapture/recovery models were fitted in JAGS (Plummer, 2003) version 4.2, using the package RJAGS version 4-6 (Plummer, 2016) in R version 3.3.2 (R Core Team, 2018). JAGS uses Markov chain Monte Carlo (MCMC) simulations to estimate parameters. We simulated three chains with 510’000 iterations, a burning phase of 10’000 iterations, and a sampling interval of 100 iterations and uninformative priors for the different parameters. The mean effective sample size for model parameter was 10’707 and the lowest effective sample size was 4’075. We visually inspected the chains and used the R-hat statistics to assess the convergence of all chains (Gelman and Hill, 2006). R-hat values for each parameter were < 1.1.

## Results

### Survival estimates

There was a clear association between baseline corticosterone levels and survival probability in male but not in female barn owls (posterior probability of baseline corticosterone levels being positively associated to survival in males (98%) and females (57%)) (Table 3, Figure 2A). In contrast, stress-induced corticosterone was associated to survival in both males and females (> 98%). Individuals presenting a higher stress-induced corticosterone level had a higher probability to survive from one year to the next (Table 3, Figure 2B). Survival probability was also associated with age in a curvilinear way (Table 3) as survival decreased in the first years of life before stabilizing at older ages. Survival was also higher in males than females, an effect that was stronger in individuals presenting a mild-to-high than low corticosterone levels (Figure 2). Finally, the probability of recapturing an individual was constant through time and hence not associated with the number of nest boxes available in the study area (Table 3).

**Figure 2.**
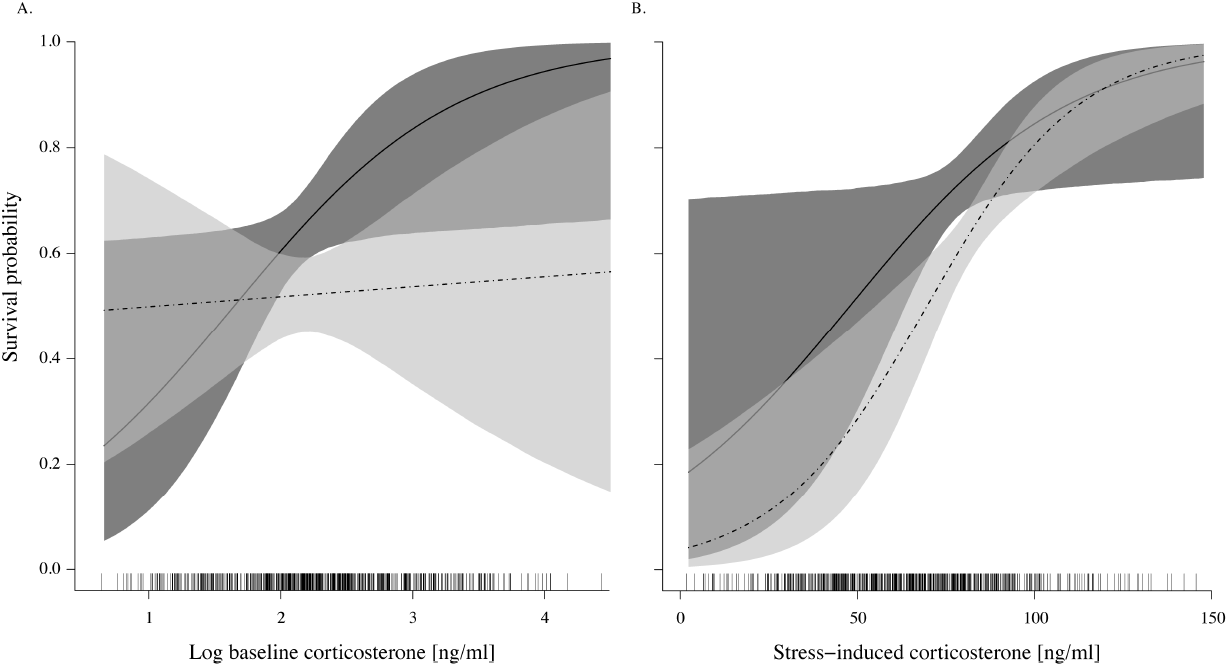
Estimated survival probability of breeding barn owls in relation to (A) baseline corticosterone levels and (B) stress-induced corticosterone levels. The black (male) and dashed lines (female) represent the mean posterior distribution estimated from (A) the baseline corticosterone model and (B) stress-induced corticosterone model (Table 3), respectively. The shaded regions represent the 95% Bayesian credible intervals. The tick marks displayed along the x-axes represent, respectively, the raw data for baseline (A) and stress-induced corticosterone levels (B).

**Table 3.**
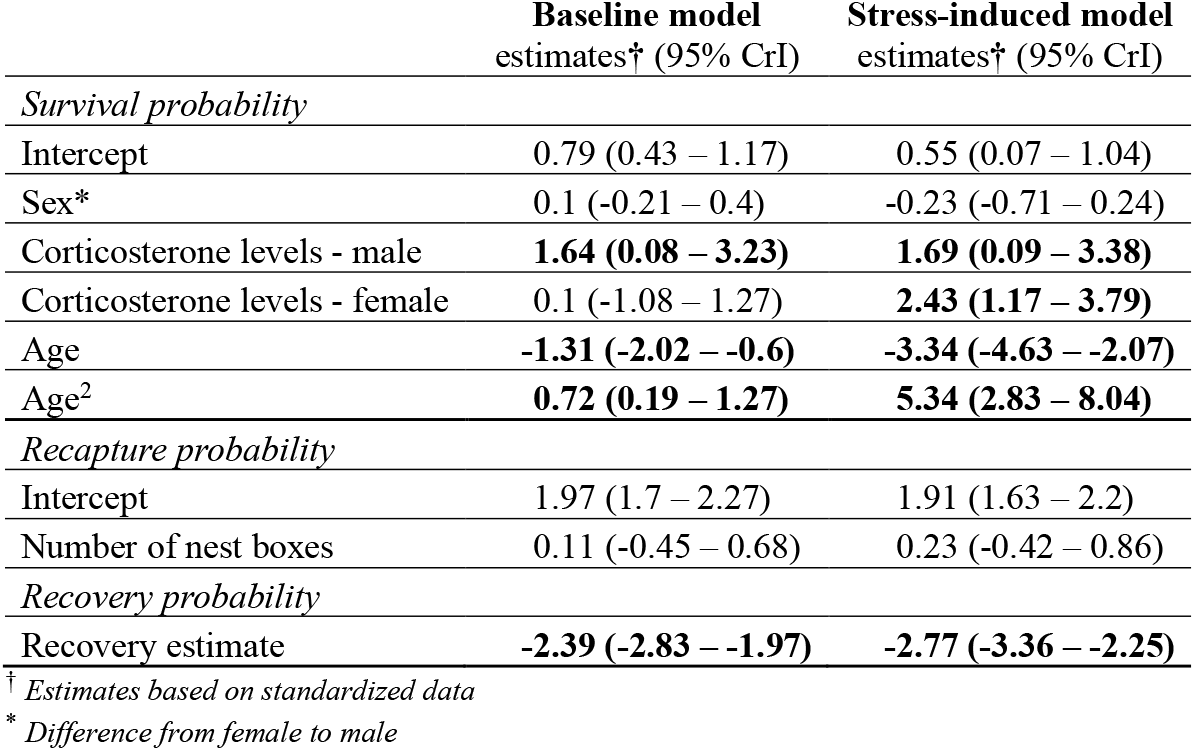
Survival estimates of breeding barn owls in relation to baseline and stress-induced corticosterone levels. Parameters are estimated from a multistate model applied to 13 years of capture-recapture data of adults monitored from 2004 to 2016. Are presented, the posterior means with 95% Bayesian credible interval for a model with only baseline (361 individuals) and stress-induced (341 individuals) corticosterone levels as predictors. Predictors with a significant effect on survival are written in bold.

## Discussion

In the present study, we examined whether variation in circulating corticosterone levels is related to survival in adult barn owls using 11 years of corticosterone measurements and 13 years of capture-recapture and dead recovery records from 2004 to 2016. Males and females presenting a higher stress-induced corticosterone response survived better than individuals showing a lower stress-induced corticosterone response (Figure 2B). We also showed that males on average have a higher survival rate than females (Figure 2) and that survival is associated with age.

### Baseline corticosterone levels and survival

Considering that glucocorticoid hormones, such as corticosterone, are essential in maintaining homeostasis and energy balance, baseline corticosterone levels are predicted to influence fitness components such as survival. For instance, high and chronically elevated corticosterone levels can induce deleterious physiological effects that can affect survival. Corticosterone has also immunosuppressive effects in different species (Bourgeon and Raclot, 2006; Fowles et al., 1993; Rubolini et al., 2005; Saino et al., 2003), including barn owl nestlings (Stier et al., 2009). Further, chronic stress is viewed as an accelerator of ageing and has been shown to reduce telomere length (e.g. Bauch et al., 2016; Monaghan, 2014), a biomarker of ageing and phenotypic quality in birds (e.g. Barrett et al., 2013; Bize et al., 2009) and humans (Boonekamp et al., 2013). Therefore, high corticosterone levels are most often predicted to be associated with low fitness (e.g. the “cort-fitness hypothesis”, Bonier et al., 2009a). Nevertheless, our results showed the opposite pattern for males, as individuals with high baseline corticosterone levels during the breeding season had a higher annual survival. Like for other fitness traits, the association between corticosterone and survival might be complex and context-dependent, as the different studies investigating the link between baseline corticosterone levels and survival have yielded mixed results, with some studies having found positive (Cabezas et al., 2007; Comendant et al., 2003; Cote et al., 2006), negative (Brown et al., 2005; Suorsa et al., 2003) or no association between baseline corticosterone levels and survival (Blas et al., 2007; Jimeno et al., 2017; Romero and Wikelski, 2001; see also Bonier et al., 2009a for review). A potential explanation for our results is that high baseline corticosterone levels could be necessary for a proper regulation of the different behavioral and physiological processes, like locomotor functions or mobilization of energetic resources (Wingfield et al., 1998) and that these individuals are in healthier condition. Elevated baseline levels may also increase the permissive action of corticosterone, enabling individuals to better respond to stressful or life-threatening situations (Sapolsky et al., 2000). For instance, some hormones involved in the stress axis, such as catecholamines and glucagon, an enzyme involved in glycogenesis, require the presence of glucocorticoids to exert their action (Exton et al., 1972; Seleznev Iu and Martynov, 1982). Individuals with high corticosterone levels might be also more prone to modify their behavior to avoid the potential cost associated with stress and thereby increase their chance of survival. For instance, adult barn owls shift their reproductive behavior towards self-maintenance (*i.e*. reduce nestling food-provisioning, home-range size and distance covered within the home-rang) under chronic stress (Almasi et al., 2013; Almasi et al., 2008). Alternatively, the low survival rate of individuals presenting low baseline corticosterone levels could be the consequence of chronic stress (Dickens and Romero, 2013), which has resulted in the down regulation and eventually the exhaustion of the HPA axis activity (Müller et al., 2009; Rich and Romero, 2005). This may have ultimately lead to a suite of deleterious health effects that have impacted their survival (Sapolsky et al., 2000).

### Stress-induced corticosterone levels and survival

Although baseline corticosterone and the acute stress response are both regulated by the HPA axis, the action and function of baseline corticosterone and the acute stress response are different, the first is mediated by the glucocorticoid type I receptor whereas the second by glucocorticoid type II (Romero, 2004). While baseline levels play a fundamental role in adjusting behavior and physiology of an organism to daily predictable changes, the acute stress response aims to maximize immediate survival during unpredictable events by suppressing temporally non-essential functions (Wingfield et al., 1998), such as reproduction or development (Meylan and Clobert, 2005; Rubolini et al., 2005; Wingfield et al., 1998). Considering that we found a positive link between stress-induced response and survival, barn owls with a higher stress-induced corticosterone response may survive better than individuals with lower stress-induced corticosterone response, as they may be able to mobilize more resources and react more adequality during an emergency response. For instance, individuals with a higher stress-induced response may show a higher fight- or flight-capacity than individuals with a lower stress-induced response (Cote et al., 2006; Overli et al., 2002a). Large increase in corticosterone is also known to stimulate activity, energy acquisition and storage through the glucocorticoid type II receptor (Dallman et al., 1995; Dallman et al., 2004; see also Vera et al., 2017 for discussion). Note that the role of the stress response is to avoid the internal system to overshoot and restore the state of homeostasis as rapidly as possible to prevent damaging effects (McEwen and Wingfield, 2003; Sapolsky et al., 2000). Therefore, an organism with large stress-induced corticosterone response might recover faster from a stress challenge as it may have a higher foraging activity (Lynn et al., 2003; Overli et al., 2002b) and food intake (Cote et al., 2006) and thereby, restore reserves more rapidly than an individual with a smaller stress-induced corticosterone response.

However, to maximize fitness, life-history theory predicts that an individual should balance costs between current reproduction and future survival (Stearns, 1989). According to this statement, individuals presenting high stress-induced corticosterone response may have a better survival but this may come with some costs that may reduce reproductive success. Indeed, individuals with a high corticosterone-response may have a reduced parental care (Lendvai and Chastel, 2010; Wingfield and Sapolsky, 2003) and reproductive success (Schmid et al., 2013; Vitousek et al., 2014). However, we adjusted corticosterone levels to parental effort (by using the mean estimated for brood size to predict baseline and stress-induced corticosterone levels) for our survival model and individuals with higher corticosterone levels still show a higher survival probability. A result that suggests that high stress responder may not trade survival against reproductive effort but that fitter individuals might just have stronger stress-response.

Corticosterone levels have also been shown to be associated to certain phenotypic traits, including personality traits (e.g. bold vs. shy) (Baugh et al., 2012; Carere et al., 2003; Pottinger and Carrick, 2001; Stowe et al., 2010), which might impact the fitness of an individual (Biro and Stamps, 2008). As a mediator of the trade-off between self-maintenance and reproduction (see review Wingfield and Sapolsky, 2003), variation in corticosterone levels may also suggest differences in life-history strategies between individuals (Lancaster et al., 2008) or coping strategies (Blas et al., 2007). This may explain why “high” and “low” stress response phenotypes are maintained in our population, in particular, if variation in stress response is associated to fitness in a context-dependent manner (Jaatinen et al., 2014).

The biological effect of corticosterone does not only depend on the total circulating levels of corticosterone (Breuner et al., 2013), but also on the density of receptors to which corticosterone binds (Seckl and Meaney, 2004), the kind and properties of receptors (Landys et al., 2006; Romero, 2004), the fraction bound to plasma corticosteroid binding globulins (Mendel, 1989), activity of enzymes involved in the metabolism of corticosterone (e.g. 11β-HSD type 1 and 2) and the negative feedback mechanism (Sapolsky et al., 2000). All these factors might also contribute for the observed variation and effects of corticosterone levels among individuals, given that all are essential in triggering the biological effects and the magnitude of such effects.

### Conclusion

Overall, our study shows that corticosterone levels are a good predictor of adult barn owl survival and that the stress-induced corticosterone response is a better predictor compared to baseline corticosterone levels (see Figure 2). This could potentially be due to the high sensitivity of baseline corticosterone levels to multiple environmental factors and life history stages in comparison to the stress-induced corticosterone response, which tends to show, in certain species, a higher stability within individuals over the years and different life history stages (Angelier et al., 2009; Cockrem et al., 2009; Hennessy et al., 2015; Rensel and Schoech, 2011; Schoenemann and Bonier, 2017). The survival advantage of individuals presenting high corticosterone levels may, however, be only advantageous at certain life history stages, in particular environments, in face of specific stressors, or for certain personalities. Again, because of the complexity and the number of factors involved in the regulation of corticosterone secretion, further studies, investigating the link between corticosterone or other components of the hypothalamic-pituitary-adrenal (HPA) axis (e.g. rapidity of feedback loop, number of glucocorticoid receptors and enzymes involves in corticosterone metabolism) and survival should be performed to examine whether corticosterone can be used as a reliable fitness proxy in birds.

## Acknowledgements

We warmly thank all field assistants for their precious help on collecting the data during the long days and nights of fieldwork. We are also grateful to the Swiss National Science Foundation who financed this research (grant n° 3100A0-104134 and 31003A-127057 to L. J. and n° 31003A-120517 to A. R.). Blood samples were taken under the legal authorization of the ‘Service vétérinaire du canton de Vaud. We are also thankful to reviewers for their comments and propositions that helped us improve the content of this paper.

## Supplementary materiel

**Table 1.** Correlation between baseline corticosterone (CORT) levels and stress-induced corticosterone levels. The estimates (95% Bayesian credible intervals) of predictors are based on the draw of 50’000 random values from the joint posterior distribution of the model parameters. The year and identity were added in the models as random terms to correct for pseudo-replication. Predictors with a significant effect on CORT levels are written in bold. The estimates are based on 518 samples taken from 356 individuals measured in 11 different years.

| Fixed effects | Stress-induced model<br>estimates<sup>†</sup> (95% CrI) |
| --- | --- |
| Intercept | 52.99 (38.93 – 67.05) |
| Baseline CORT | **16.89 (11.98 – 21.85)** |
| Baseline CORT<sup>2</sup> | **-6.85 (-11.57 – -2.16)** |
| Date of sampling | **-3.35 (-5.19 – -1.54)** |
| Hour of sampling | 1.28 (-0.46 – 3.01) |
| Blood sampling time* | 1.7 (-0.79 – 4.16) |
| Body condition | **-6.59 (-8.79 – -4.37)** |
| Sex | 6.19 (-0.2 – 12.51) |

<sup>†</sup> Estimates based on standardized data
\* latency between time (sec) of capture and blood sampling for stress-induced model.

## References

Almasi, B., Béziers, P., Roulin, A., Jenni, L., 2015. Agricultural land use and human presence around breeding sites increase stress-hormone levels and decrease body mass in barn owl nestlings. Oecologia 179, 89–101.

Almasi, B., Roulin, A., Jenni, L., 2013. Corticosterone shifts reproductive behaviour towards self-maintenance in the barn owl and is linked to melanin-based coloration in females. Horm Behav 64, 161–171.

Almasi, B., Roulin, A., Jenni-Eiermann, S., Jenni, L., 2008. Parental investment and its sensitivity to corticosterone is linked to melanin-based coloration in barn owls. Horm Behav 54, 217–223.

Altwegg, R., Schaub, M., Roulin, A., 2007. Age-specific fitness components and their temporal variation in the barn owl. Am Nat 169, 47–61.

Angelier, F., Ballentine, B., Holberton, R.L., Marra, P.P., Greenberg, R., 2011. What drives variation in the corticosterone stress response between subspecies? A common garden experiment of swamp sparrows (Melospiza georgiana). J Evolution Biol 24, 1274–1283.

Angelier, F., Holberton, R.L., Marra, P.P., 2009. Does stress response predict return rate in a migratory bird species? A study of American redstarts and their non-breeding habitat. P R Soc B 276, 3545–3551.

Barrett, E.L.B., Burke, T.A., Hammers, M., Komdeur, J., Richardson, D.S., 2013. Telomere length and dynamics predict mortality in a wild longitudinal study. Mol Ecol 22, 249–259.

Bates, D., Machler, M., Bolker, B.M., Walker, S.C., 2015. Fitting Linear Mixed-Effects Models Using lme4. J Stat Softw 67, 1–48.

Bauch, C., Riechert, J., Verhulst, S., Becker, P.H., 2016. Telomere length reflects reproductive effort indicated by corticosterone levels in a long-lived seabird. Mol Ecol 25, 5785–5794.

Baugh, A.T., Schaper, S.V., Hau, M., Cockrem, J.F., de Goede, P., van Oers, K., 2012. Corticosterone responses differ between lines of great tits (Parus major) selected for divergent personalities. Gen Comp Endocr 175, 488–494.

Béziers, P., Roulin, A., 2016. Double brooding and offspring desertion in the barn owl Tyto alba. J Avian Biol 47, 235–244.

Biro, P.A., Stamps, J.A., 2008. Are animal personality traits linked to life-history productivity? Trends Ecol Evol 23, 361–368.

Bize, P., Criscuolo, F., Metcalfe, N.B., Nasir, L., Monaghan, P., 2009. Telomere dynamics rather than age predict life expectancy in the wild. P R Soc B 276, 1679–1683.

Bize, P., Stocker, A., Jenni-Eiermann, S., Gasparini, J., Roulin, A., 2010. Sudden weather deterioration but not brood size affects baseline corticosterone levels in nestling Alpine swifts. Horm Behav 58, 591–598.

Blas, J., Bortolotti, G.R., Tella, J.L., Baos, R., Marchant, T.A., 2007. Stress response during development predicts fitness in a wild, long lived vertebrate. Proceedings of the National Academy of Sciences of the United States of America 104, 8880–8884.

Bonier, F., Martin, P.R., 2016. How can we estimate natural selection on endocrine traits? Lessons from evolutionary biology. P R Soc B 283.

Bonier, F., Martin, P.R., Moore, I.T., Wingfield, J.C., 2009a. Do baseline glucocorticoids predict fitness? Trends Ecol Evol 24, 634–642.

Bonier, F., Martin, P.R., Sheldon, K.S., Jensen, J.P., Foltz, S.L., Wingfield, J.C., 2007. Sex-specific consequences of life in the city. Behav Ecol 18, 121–129.

Bonier, F., Moore, I.T., Martin, P.R., Robertson, R.J., 2009b. The relationship between fitness and baseline glucocorticoids in a passerine bird. Gen Comp Endocr 163, 208–213.

Boonekamp, J.J., Simons, M.J.P., Hemerik, L., Verhulst, S., 2013. Telomere length behaves as biomarker of somatic redundancy rather than biological age. Aging Cell 12, 330–332.

Bourgeon, S., Raclot, T., 2006. Corticosterone selectively decreases humoral immunity in female eiders during incubation. J Exp Biol 209, 4957–4965.

Bremner, J.D., 2007. Does Stress Damage the Brain? Understanding Trauma: Integrating Biological, Clinical, and Cultural Perspectives, 118–141.

Breuner, C.W., Delehanty, B., Boonstra, R., 2013. Evaluating stress in natural populations of vertebrates: total CORT is not good enough. Funct Ecol 27, 24–36.

Breuner, C.W., Patterson, S.H., Hahn, T.P., 2008. In search of relationships between the acute adrenocortical response and fitness. Gen Comp Endocr 157, 288–295.

Brown, C.R., Brown, M.B., Raouf, S.A., Smith, L.C., Wingfield, J.C., 2005. Effects of endogenous steroid hormone levels on annual survival in Cliff Swallows. Ecology 86, 1034–1046.

Brown, K.I., Nestor, K.E., 1973. Some Physiological Responses of Turkeys Selected for High and Low Adrenal Response to Cold Stress. Poultry Science 52, 1948–1954.

Cabezas, S., Blas, J., Marchant, T.A., Moreno, S., 2007. Physiological stress levels predict survival probabilities in wild rabbits. Horm Behav 51, 313–320.

Carere, C., Groothuis, T.G.G., Mostl, E., Daan, S., Koolhaas, J.M., 2003. Fecal corticosteroids in a territorial bird selected for different personalities: daily rhythm and the response to social stress. Horm Behav 43, 540–548.

Carsia, R.V., Harvey, A., 2000. Adrenals, in: Causey, G.W. (Ed.), Sturkie’s Avian Physiology, 5th ed. Academic Press, London, pp. 489–522.

Cockrem, J.F., Barrett, D.P., Candy, E.J., Potter, M.A., 2009. Corticosterone responses in birds: Individual variation and repeatability in Adelie penguins (Pygoscelis adeliae) and other species, and the use of power analysis to determine sample sizes. Gen Comp Endocr 163, 158–168.

Comendant, T., Sinervo, B., Svensson, E.I., Wingfield, J., 2003. Social competition, corticosterone and survival in female lizard morphs. J Evolution Biol 16, 948–955.

Cote, J., Clobert, J., Meylan, S., Fitze, P.S., 2006. Experimental enhancement of corticosterone levels positively affects subsequent male survival. Horm Behav 49, 320–327.

Crespi, E.J., Williams, T.D., Jessop, T.S., Delehanty, B., 2013. Life history and the ecology of stress: how do glucocorticoid hormones influence life-history variation in animals? Funct Ecol 27, 93–106.

Crossin, G.T., Love, O.P., Cooke, S.J., Williams, T.D., 2016. Glucocorticoid manipulations in free-living animals: considerations of dose delivery, life-history context and reproductive state. Funct Ecol 30, 116–125.

Dallman, M.F., Akana, S.F., Strack, A.M., Hanson, E.S., Sebastian, R.J., 1995. The neural network that regulates energy balance is responsive to glucocorticoids and insulin and also regulates HPA axis responsivity at a site proximal to CRF neurons. Stress 771, 730–742.

Dallman, M.F., La Fleur, S.E., Pecoraro, N.C., Gomez, F., Houshyar, H., Akana, S.F., 2004. Minireview: Glucocorticoids - Food intake, abdominal obesity, and wealthy nations in 2004. Endocrinology 145, 2633–2638.

Dallman, M.F., Strack, A.M., Akana, S.F., Bradbury, M.J., Hanson, E.S., Scribner, K.A., Smith, M., 1993. Feast and Famine - Critical Role of Glucocorticoids with Insulin in Daily Energy-Flow. Front Neuroendocrin 14, 303–347.

Dantzer, B., Newman, A.E.M., Boonstra, R., Palme, R., Boutin, S., Humphries, M.M., McAdam, A.G., 2013. Density Triggers Maternal Hormones That Increase Adaptive Offspring Growth in a Wild Mammal. Science 340, 1215–1217.

de Kloet, E.R., Oitzl, M.S., Joels, M., 1999. Stress and cognition: are corticosteroids good or bad guys? Trends Neurosci 22, 422–426.

Dickens, M.J., Romero, L.M., 2013. A consensus endocrine profile for chronically stressed wild animals does not exist. Gen Comp Endocr 191, 177–189.

Dreiss, A.N., Roulin, A., 2014. Divorce in the barn owl: securing a compatible or better mate entails the cost of re-pairing with a less ornamented female mate. J Evolution Biol 27, 1114–1124.

Dunn, A.J., Powell, M.L., Meitin, C., Small, P.A., 1989. Virus-Infection as a Stressor - Influenza-Virus Elevates Plasma-Concentrations of Corticosterone, and Brain Concentrations of Mhpg and Tryptophan. Physiol Behav 45, 591–594.

Evans, M.R., Roberts, M.L., Buchanan, K.L., Goldsmith, A.R., 2006. Heritability of corticosterone response and changes in life history traits during selection in the zebra finch. J Evol Biol 19, 343–352.

Exton, J.H., Friedmann, N., Wong, E.H.A., Brineaux, J.P., Corbin, J.D., Park, C.R., 1972. Interaction of glucocorticoids with glucagon and epinephrine in control of gluconeogenesis and glycogenolysis in liver and of lipolysis in adipose-tissue. J Biol Chem 247, 3579-+.

Fowles, J.R., Fairbrother, A., Fix, M., Schiller, S., Kerkvliet, N.I., 1993. Glucocorticoid effects on natural and humoral immunity in mallards. Developmental and Comparative Immunology 17, 165–177.

Gelman, A., Hill, J., 2006. Data Analysis Using Regression and Multilevel/Hierarchical Models. Cambridge University Press, NewYork.

Glennemeier, K.A., Denver, R.J., 2002. Role for corticoids in mediating the response of Rana pipiens tadpoles to intraspecific competition. J Exp Zool 292, 32–40.

Gustafsson, L., Nordling, D., Andersson, M.S., Sheldon, B.C., Qvarnstrom, A., 1994. Infectious-Diseases, Reproductive Effort and the Cost of Reproduction in Birds. Philos T Roy Soc B 346, 323–331.

Hau, M., Ricklefs, R.E., Wikelski, M., Lee, K.A., Brawn, J.D., 2010. Corticosterone, testosterone and life-history strategies of birds. P R Soc B 277, 3203–3212.

Hennessy, M.B., Kaiser, S., Tiedtke, T., Sachser, N., 2015. Stability and change: Stress responses and the shaping of behavioral phenotypes over the life span. Front Zool 12.

Jaatinen, K., Seltmann, M.W., Ost, M., 2014. Context-dependent stress responses and their connections to fitness in a landscape of fear. J Zool 294, 147–153.

Jenkins, B.R., Vitousek, M.N., Hubbard, J.K., Safran, R.J., 2014. An experimental analysis of the heritability of variation in glucocorticoid concentrations in a wild avian population. P R Soc B 281.

Jenni-Eiermann, S., Glaus, E., Gruebler, M., Schwabl, H., Jenni, L., 2008. Glucocorticoid response to food availability in breeding barn swallows (Hirundo rustica). Gen Comp Endocr 155, 558–565.

Jimeno, B., Michael, B., Hau, M., Verhulst, S., 2017. Male but not female zebra finches with high plasma corticosterone have lower survival. Funct Ecol.

King, B.M., 1988. Glucocorticoids and Hypothalamic Obesity. Neurosci Biobehav R 12, 29–37.

Kitaysky, A.S., Piatt, J.F., Wingfield, J.C., 2007. Stress hormones link food availability and population processes in seabirds. Mar Ecol Prog Ser 352, 245–258.

Lancaster, L.T., Hazard, L.C., Clobert, J., Sinervo, B.R., 2008. Corticosterone manipulation reveals differences in hierarchical organization of multidimensional reproductive trade-offs in r-strategist and K-strategist females. J Evolution Biol 21, 556–565.

Landys, M.M., Ramenofsky, M., Wingfield, J.C., 2006. Actions of glucocorticoids at a seasonal baseline as compared to stress-related levels in the regulation of periodic life processes. Gen Comp Endocr 148, 132–149.

Lebreton, J.D., Nichols, J.D., Barker, R.J., Pradel, R., Spendelow, J.A., 2009. Modeling Individual Animal Histories with Multistate Capture-Recapture Models, in: Caswell, H. (Ed.), Advances in Ecological Research, Vol 41, pp. 87–173.

Lendvai, A.Z., Chastel, O., 2010. Natural variation in stress response is related to post-stress parental effort in male house sparrows. Horm Behav 58, 936–942.

Little, R.J.A., Rubin, D.B., 2002. Statistical analysis with missing data, second edition, New York, NY.

Love, A.C., Foltz, S.L., Adelman, J.S., Moore, I.T., Hawley, D.M., 2016. Changes in corticosterone concentrations and behavior during Mycoplasma gallisepticum infection in house finches (Haemorhous mexicanus). Gen Comp Endocr 235, 70–77.

Lynn, S.E., Breuner, C.W., Wingfield, J.C., 2003. Short-term fasting affects locomotor activity, corticosterone, and corticosterone binding globulin in a migratory songbird. Horm Behav 43, 150–157.

Magee, S.E., Neff, B.D., Knapp, R., 2006. Plasma levels of androgens and cortisol in relation to breeding behavior in parental male bluegill sunfish, Lepomis macrochirus. Horm Behav 49, 598–609.

Martin, L.B., 2009. Stress and immunity in wild vertebrates: Timing is everything. Gen Comp Endocr 163, 70–76.

McEwen, B.S., Wingfield, J.C., 2003. The concept of allostasis in biology and biomedicine. Horm Behav 43, 2–15.

Mendel, C.M., 1989. The free hormone hypothesis: a physiologically based mathematical model. Endocr Rev 10, 232–274.

Meylan, S., Clobert, J., 2004. Maternal effects on offspring locomotion: Influence of density and corticosterone elevation in the lizard Lacerta vivipara. Physiol Biochem Zool 77, 450–458.

Meylan, S., Clobert, J., 2005. Is corticosterone-mediated phenotype development adaptive? - Maternal corticosterone treatment enhances survival in male lizards. Horm Behav 48, 44–52.

Monaghan, P., 2014. Organismal stress, telomeres and life histories. J Exp Biol 217, 57–66.

Müller, C., Almasi, B., Roulin, A., Breuner, C.W., Jenni-Eiermann, S., Jenni, L., 2009. Effects of corticosterone pellets on baseline and stress-induced corticosterone and corticosteroid-binding-globulin. Gen Comp Endocr 160, 59–66.

Müller, C., Jenni-Eiermann, S., Blondel, J., Perret, P., Caro, S.P., Lambrechts, M., Jenni, L., 2006. Effect of human presence and handling on circulating corticosterone levels in breeding blue tits (Parus caeruleus). Gen Comp Endocr 148, 163–171.

Munro, C., Stabenfeldt, G., 1984. Development of a microtitre plate enzyme immunoassay for the determination of progesterone. J Endocrinol 101, 41–49.

Munro, C.J., Lasley, B.L., 1988. Non-radiometric methods for immunoassay of steroid hormones. Progress in clinical and biological research 285, 289–329.

Odeh, F.M., Cadd, G.G., Satterlee, D.G., 2003. Genetic characterization of stress responsiveness in Japanese quail. 2. Analyses of maternal effects, additive sex linkage effects, heterosis, and heritability by diallel crosses. Poultry Science 82, 31–35.

Ouyang, J.Q., Lendvai, A.Z., Dakin, R., Domalik, A.D., Fasanello, V.J., Vassallo, B.G., Haussmann, M.F., Moore, I.T., Bonier, F., 2015. Weathering the storm: parental effort and experimental manipulation of stress hormones predict brood survival. Bmc Evol Biol 15.

Ouyang, J.Q., Quetting, M., Hau, M., 2012. Corticosterone and brood abandonment in a passerine bird. Anim Behav 84, 261–268.

Overli, O., Kotzian, S., Winberg, S., 2002a. Effects of cortisol on aggression and locomotor activity in rainbow trout. Horm Behav 42, 53–61.

Overli, O., Pottinger, T.G., Carrick, T.R., Overli, E., Winberg, S., 2002b. Differences in behaviour between rainbow trout selected for high- and low-stress responsiveness. J Exp Biol 205, 391–395.

Patterson, S.H., Hahn, T.P., Cornelius, J.M., Breuner, C.W., 2014. Natural selection and glucocorticoid physiology. J Evolution Biol 27, 259–274.

Plummer, M., 2003. JAGS: a program for analysis of Bayesiangraphical models using Gibbs sampling, Proceedings of the 3rdInternational Workshop on Distributed Statistical Computing, Vienna, Austria.

Plummer, M., 2016. rjags: Bayesian Graphical Models usingMCMC. R package version 4–6.

Pottinger, T.G., Carrick, T.R., 1999. Modification of the plasma cortisol response to stress in rainbow trout by selective breeding. Gen Comp Endocrinol 116, 122–132.

Pottinger, T.G., Carrick, T.R., 2001. Stress responsiveness affects dominant-subordinate relationships in rainbow trout. Horm Behav 40, 419–427.

Rensel, M.A., Schoech, S.J., 2011. Repeatability of baseline and stress-induced corticosterone levels across early life stages in the Florida scrub-jay (Aphelocoma coerulescens). Horm Behav 59, 497–502.

Rich, E.L., Romero, L.M., 2005. Exposure to chronic stress downregulates corticosterone responses to acute stressors. Am J Physiol-Reg I 288, R1628–R1636.

Ricklefs, R.E., Wikelski, M., 2002. The physiology/life-history nexus. Trends Ecol Evol 17, 462–468.

Riechert, J., Becker, P.H., Chastel, O., 2014. Predicting reproductive success from hormone concentrations in the common tern (Sterna hirundo) while considering food abundance. Oecologia 176, 715–727.

Rivers, J.W., Liebl, A.L., Owen, J.C., Martin, L.B., Betts, M.G., 2012. Baseline corticosterone is positively related to juvenile survival in a migrant passerine bird. Funct Ecol 26, 1127–1134.

Romero, L.M., 2002. Seasonal changes in plasma glucocorticoid concentrations in free-living vertebrates. Gen Comp Endocr 128, 1–24.

Romero, L.M., 2004. Physiological stress in ecology: lessons from biomedical research. Trends Ecol Evol 19, 249–255.

Romero, L.M., Reed, J.M., 2005. Collecting baseline corticosterone samples in the field: is under 3 min good enough? Comp Biochem Phys A 140, 73–79.

Romero, L.M., Wikelski, M., 2001. Corticosterone levels predict survival probabilities of Galapagos marine iguanas during El Nino events. Proceedings of the National Academy of Sciences of the United States of America 98, 7366–7370.

Roulin, A., Almasi, B., Jenni, L., 2010. Temporal variation in glucocorticoid levels during the resting phase is associated in opposite way with maternal and paternal melanic coloration. J Evolution Biol 23, 2046–2053.

Rubolini, D., Romano, M., Boncoraglio, G., Ferrari, R.P., Martinelli, R., Galeotti, P., Fasola, M., Saino, N., 2005. Effects of elevated egg corticosterone levels on behavior, growth, and immunity of yellow-legged gull (Larus michahellis) chicks. Horm Behav 47, 592–605.

Saino, N., Suffritti, C., Martinelli, R., Rubolini, D., Moller, A.P., 2003. Immune response covaries with corticosterone plasma levels under experimentally stressful conditions in nestling barn swallows (Hirundo rustica). Behav Ecol 14, 318–325.

Sapolsky, R.M., Romero, L.M., Munck, A.U., 2000. How do glucocorticoids influence stress responses? Integrating permissive, suppressive, stimulatory, and preparative actions. Endocr Rev 21, 55–89.

Satterlee, D.G., Johnson, W.A., 1988. Selection of Japanese quail for contrasting blood corticosterone response to immobilization. Poult Sci 67, 25–32.

Scheuerlein, A., Van’t Hof, T.J., Gwinner, E., 2001. Predators as stressors? Physiological and reproductive consequences of predation risk in tropical stonechats (Saxicola torquata axillaris). P R Soc B 268, 1575–1582.

Schmid, B., Tam-Dafond, L., Jenni-Eiermann, S., Arlettaz, R., Schaub, M., Jenni, L., 2013. Modulation of the adrenocortical response to acute stress with respect to brood value, reproductive success and survival in the Eurasian hoopoe. Oecologia 173, 33–44.

Schoenemann, K.L., Bonier, F., 2017. Repeatability of glucocorticoid hormones in vertebrates: A meta-analysis. Peerj.

Schultner, J., Kitaysky, A.S., Gabrielsen, G.W., Hatch, S.A., Bech, C., 2013. Differential reproductive responses to stress reveal the role of life-history strategies within a species. P R Soc B 280.

Seckl, J.R., Meaney, M.J., 2004. Glucocorticoid programming. Biobehavioral Stress Response: Protective and Damaging Effects 1032, 63–84.

Seleznev Iu, M., Martynov, A.V., 1982. Permissive effect of glucocorticoids in catecholamine action in the heart: possible mechanism. J Mol Cell Cardiol 14 Suppl 3, 49–58.

Sheriff, M.J., Krebs, C.J., Boonstra, R., 2009. The sensitive hare: sublethal effects of predator stress on reproduction in snowshoe hares. J Anim Ecol 78, 1249–1258.

Sopinka, N.M., Patterson, L.D., Redfern, J.C., Pleizier, N.K., Belanger, C.B., Midwood, J.D., Crossin, G.T., Cooke, S.J., 2015. Manipulating glucocorticoids in wild animals: basic and applied perspectives. Conserv Physiol 3.

Stier, K.S., Almasi, B., Gasparini, J., Piault, R., Roulin, A., Jenni, L., 2009. Effects of corticosterone on innate and humoral immune functions and oxidative stress in barn owl nestlings. J Exp Biol 212, 2084–2090.

Stowe, M., Rosivall, B., Drent, P.J., Mostl, E., 2010. Selection for fast and slow exploration affects baseline and stress-induced corticosterone excretion in Great tit nestlings, Parus major. Horm Behav 58, 864–871.

Suorsa, P., Huhta, E., Nikula, A., Nikinmaa, M., Jantti, A., Helle, H., Hakkarainen, H., 2003. Forest management is associated with physiological stress in an old-growth forest passerine. P R Soc B 270, 963–969.

Taylor, I.R., 1993. Age and sex determination of barn owls Tyto alba alba. Ringing and Migration 14, 94–102.

Thierry, A.M., Massemin, S., Handrich, Y., Raclot, T., 2013. Elevated corticosterone levels and severe weather conditions decrease parental investment of incubating Adelie penguins. Horm Behav 63, 475–483.

Vera, F., Zenuto, R., Antenucci, C.D., 2017. Expanding the actions of cortisol and corticosterone in wild vertebrates: A necessary step to overcome the emerging challenges. Gen Comp Endocr 246, 337–353.

Vitousek, M.N., Jenkins, B.R., Safran, R.J., 2014. Stress and success: Individual differences in the glucocorticoid stress response predict behavior and reproductive success under high predation risk. Horm Behav 66, 812–819.

Wey, T.W., Lin, L., Patton, M.L., Blumstein, D.T., 2015. Stress hormone metabolites predict overwinter survival in yellow-bellied marmots. Acta Ethol 18, 181–185.

Wilkening, J.L., Ray, C., 2016. Characterizing predictors of survival in the American pika (Ochotona princeps). J Mammal 97, 1366–1375.

Wingfield, J.C., Maney, D.L., Breuner, C.W., Jacobs, J.D., Lynn, S., Ramenofsky, M., Richardson, R.D., 1998. Ecological bases of hormone-behavior interactions: The “emergency life history stage”. Am Zool 38, 191–206.

Wingfield, J.C., Sapolsky, R.M., 2003. Reproduction and resistance to stress: When and how. J Neuroendocrinol 15, 711–724.

Zera, A.J., Harshman, L.G., Williams, T.D., 2007. Evolutionary endocrinology: The developing synthesis between endocrinology and evolutionary genetics. Annu Rev Ecol Evol S 38, 793–817.

